# Differential Immunomodulatory Properties of Langya and Nipah Virus Proteins

**DOI:** 10.1101/2025.09.09.675060

**Authors:** Stéphanie Durand, Dimitrios Marousis, Branka Horvat, Louis-Marie Bloyet

## Abstract

Langya virus (LayV) is a shrew-borne emerging parahenipavirus first identified in 2018 in 35 febrile patients in China. The closely related Nipah virus (NiV) is a highly pathogenic emerging bat-borne henipavirus that has caused numerous outbreaks with public health concerns in Asia. Among other symptoms, NiV causes severe acute respiratory syndrome and encephalitis, leading to high lethality. Thus, although closely related, infections with these two emerging and zoonotic viruses have distinct pathogenicity. Since the interplay with the host’s immune system is a key determinant of species barrier crossing and pathogenicity, we aimed at deciphering the ability of LayV to counteract the human intrinsic immunity using the better-characterised NiV as a prototype. NiV expresses the P, V, and W proteins, known to hinder the host’s innate immune response during infection. We thus compared the immunomodulatory properties of LayV and NiV proteins in human cells and showed that, similarly to NiV, the C-terminal domain of LayV V proteins inhibits the response to the activation of the pattern recognition receptor MDA5. However, although the N-terminal region of LayV P can inhibit the interferon signalling, it is not as efficient as its NiV counterpart. Moreover, only NiV W inhibits MDA5 and RIG-I signalling pathways. Similarly, unlike NiV, LayV W cannot efficiently block the activation of the NF-κB promoter after stimulation with IL-1β. These results suggest that LayV is less efficient than NiV in counteracting the human intrinsic immunity, which may contribute to the difference in severity observed between NiV and LayV-infected patients.

**IMPORTANCE:** Langya virus (LayV) is a shrew-borne emerging parahenipavirus recently identified in patients in China with symptoms such as fever, fatigue, cough, anorexia, headache, and vomiting. Since the interaction between a virus and its host’s immune system is an essential parameter influencing host adaptation and disease severity, we investigated the interplay between LayV and the human intrinsic immunity using as a prototype the better-characterized and closely related Nipah virus (NiV), a highly pathogenic henipavirus. We compared the immunomodulatory properties of LayV and NiV proteins in human cells and showed that, similarly to NiV, some of LayV proteins can inhibit human signalling pathways, while, unlike NiV, LayV W protein is unable to block essential immune pathways. This suggests that LayV is less efficient than NiV in counteracting the human intrinsic immunity, which may contribute to the difference in severity between NiV and LayV-infected patients.

## INTRODUCTION

Langya and Nipah viruses are closely related zoonotic paramyxoviruses responsible for outbreaks in Asia. Langya virus (LayV) was first identified in 2018 in 35 patients with acute infections in China’s Henan and Shandong provinces [1]. Symptoms included fever, fatigue, cough, anorexia, myalgia, nausea, headache, and vomiting. Shrews are likely the natural reservoir of LayV, although seropositivity has also been detected in dogs and goats. LayV has also been recently identified in shrews in the Republic of Korea [2], and closely related viruses have been discovered in shrews and rodents in Asia, Africa, Europe, and North America [3–10]. Due to its phylogenetic proximity with Nipah and Hendra viruses, LayV was initially described as a henipavirus, but the discovery of several related shrew-borne viruses led to the creation of the new *Parahenipavirus* genus [11] (Figure 1A). The ability of LayV to infect humans, the severity of the symptoms, the seropositivity of domestic animals, and its widespread geographic area raise concerns about its zoonotic potential. To date, no reverse genetic system for LayV is available, only the structures of LayV glycoproteins have been studied [12–17], and the interplay between LayV and its hosts remains unexplored.

**Figure 1.**
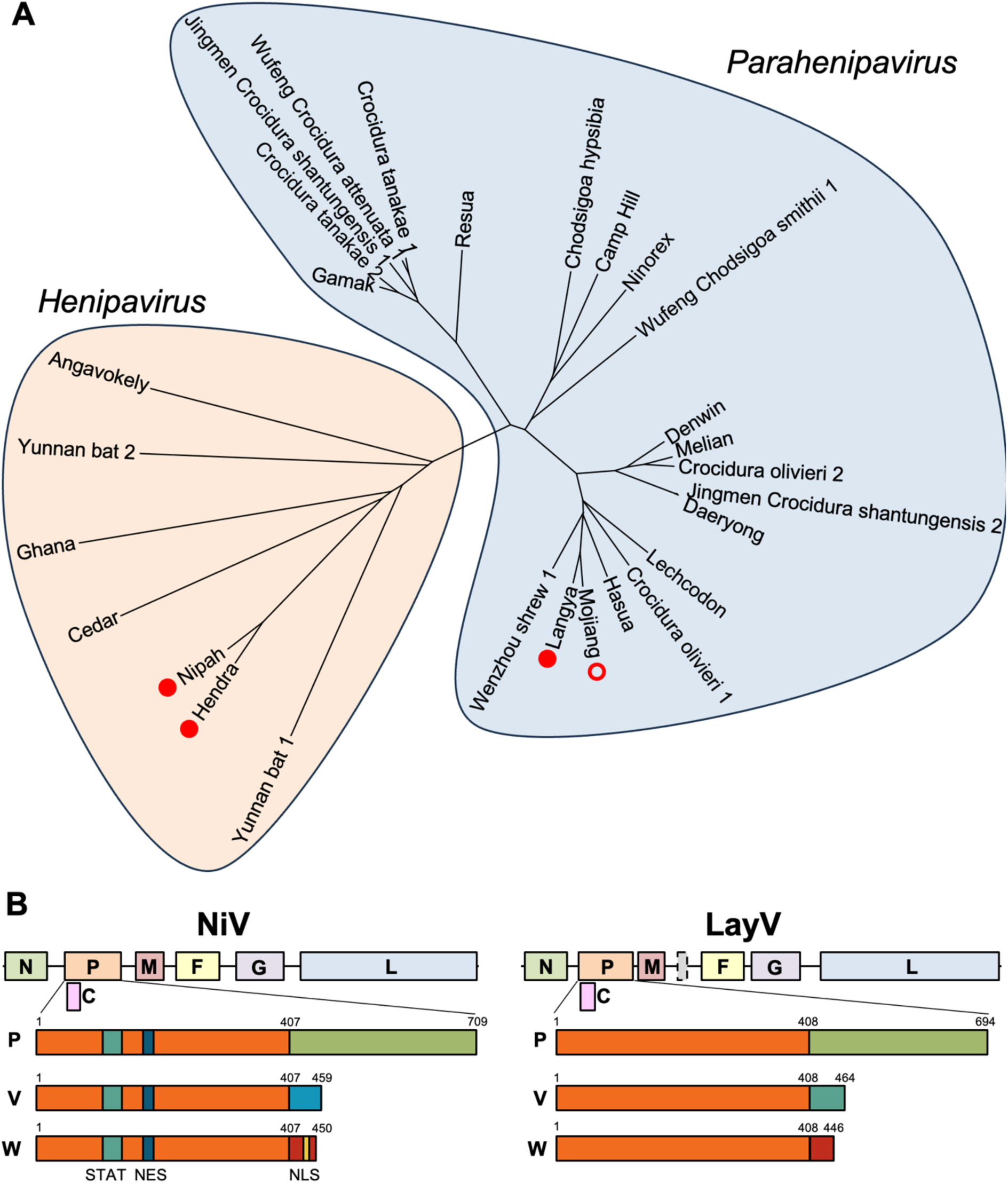
Phylogenetic tree and genome organisation of henipaviruses and parahenipaviruses. (A) Phylogenetic tree of henipavirus and parahenipaviruses based on the protein sequence of the L protein. Red dots indicate viruses known to be pathogenic in humans, and the red circle indicates Mojiang virus, suspected to be pathogenic for humans [5]. (B) Genome organisation of NiV and LayV. The proteins produced from the P gene are indicated below the genomes. A STAT1-binding site (STAT), a nucleus export signal (NES), and a nucleus localisation signal (NLS), described for NiV, are indicated.

Nipah virus (NiV) is a closely related emerging and highly pathogenic virus responsible for regular zoonotic outbreaks in Asia [18]. Discovered during an outbreak in Malaysia in 1998, NiV causes severe symptoms, including severe acute respiratory syndrome and encephalitis, with a case-fatality rate ranging from 40% to 75%. Since 2001, almost annual outbreaks in Bangladesh and India have been caused by spillovers from NiV natural reservoir: *Pteropus* fruit bats. In the absence of approved vaccines and treatments, and reports of human-to-human transmission, the World Health Organisation placed NiV on its Blueprint list of priority pathogens requiring urgent research and development efforts [19].

NiV and LayV share a similar genome organisation with the conserved six adjacent genes (Figure 1B) [2]. Most henipavirus P genes encode four proteins: P, V, W, and C proteins [20]. V and W proteins are expressed from mRNAs edited by the viral polymerase during transcription with the addition of one (V) or two (W) guanosine residues at a specific editing site. Thus, the P, V, and W proteins share a common N-terminal domain (P_NTD_) but differ in their unique C-terminal domains (CTD) (Figure 1B). Although there is no direct evidence for the existence of LayV V and W proteins, the conservation of the editing site and the sequences of the V and W CTDs strongly suggests these proteins are produced during the infection (Figure S1B-D).

NiV is well known to interfere with the host’s intrinsic immunity, notably by inhibiting the interferon type I (IFN-I) induction and signalling pathways, contributing to its high pathogenicity [21–26]. Indeed, P_NTD_, common to P, V, and W, binds and inhibits STAT1, thus preventing the expression of interferon-stimulated genes (ISGs) [27–31,26,32,33]. Additionally, NiV V protein binds to MDA5, disrupts its folding, and thus prevents its activation and the expression of interferon beta (IFN-β) [34–36]. Via its CTD, NiV W also inhibits the NF-κB pathway by preventing the accumulation of the p65 subunit in the nucleus [37].

Although closely related, LayV and NiV have different pathogenicity and infection outcomes in humans. Since the interplay between viruses and the innate immune response plays a critical role in pathogenicity, to examine these differences we compared the ability of the P, V, and W proteins of LayV and NiV to counteract several pathways of the intrinsic immunity in human cells.

## MATERIALS AND METHODS

### Cells

Human carcinoma HeLa cell line (Cat# ATCC® CCL-2™), Human Embryonic Kidney HEK293T (Cat# ATCC® CRL-3216™) and Human hepatocellular carcinoma cell line Huh-7.5 (kind gift from Charles Rice (Rockefeller University, New York, USA)) were grown in Dulbecco’s modified Eagle’s medium (DMEM), high glucose, GlutaMAX™ (LifeTechnologies™, Cat# 61965-026) supplemented with 10% of heat-inactivated (30 min at 56°C) foetal calf serum (Gibco, Cat #A5256701), and 1% of Penicillin-Streptomycin solution (Life Technologies™, Cat# 15140-130). All cell types were incubated at 37°C with 5% CO_2_ and tested negative for Mycoplasma *spp*. (Lonza, Cat#LT07-318). Depending on their features, the different cell lines were used in specific assays: since they have a very high transfection rate, HEK293T cells were used in the IFN-I response luciferase assay to maximize the signal-to-noise ratio; due to their large cytoplasm, HeLa cells were used for microscopy and, to be consistent with the p65 localization assay, for the NF-κB pathway luciferase assay; and Huh7.5 cells were used to study RIG-I and MDA5 since they do not express RIG-I and only a low level of MDA5, thus allowing us to study one sensor or the other without interference from endogenous proteins [38].

### Plasmids

The plasmids expressing the Firefly luciferase under the control of the IFN-β promoter (pβ-IFN-Luc [39]), the NF-κB promoter (pSGNluc, Addgene, Cat# 186705) [40], and the interferon response element (pISRE-Luc, Stratagene, Cat # 219092) were used to monitor the activation of the promoters. The plasmid phRL-TK expressing the Renilla luciferase under the HSV-thymidine kinase promoter (Promega, Cat# E6241) was used for normalisation. The plasmids pEF_c-myc-MDA5 and pEF-BOS-HA-RIG-I encode myc-tagged human MDA5 and HA-tagged human RIG-I, respectively [41].

Plasmids coding for LayV P, V, and W (GenBank #OM101129) were cloned in pcDNA3.1 and ordered from GeneScript. These proteins contain an N-terminal Flag tag, separated by a GGGSG linker. Plasmids coding for NiV proteins (GenBank #AY988601) were cloned with the same tag in the same backbone using InFusion-mediated recombination (Takara, Cat# 638933). To express P/V/W_NTD_, P_CTD_ was removed from the plasmids expressing P. V_CTD_ and W_CTD_ were expressed as fusion proteins with a Flag-tagged mCherry protein fused at their N-terminus (separated from the CTDs by a single G residue). V and W mutants were generated from the plasmids encoding the non-mutated proteins using InFusion-mediated recombination.

The open reading frames of all plasmids were sequenced by Sanger sequencing (Eurofins Genomics).

### Luciferase reporter assays

For the investigation of the MDA5 and RIG-I pathways, 2×10^4^ Huh-7.5 cells were seeded per well in ninety-six-well plates. Each well was transfected the day after with 50 ng of pβ-IFN-Luc, 20 ng of phRL-TK, 50 ng of MDA5 or RIG-I expressing plasmid, together with 80 ng of plasmids expressing the viral proteins or the mCherry control, using JetOptimus reagent (Ozyme, Cat# POL101000006) according to the manufacturer’s recommendations.

Cells were transfected 24 h later with 100 ng of Poly(I:C) (Invivogen, Cat# PIC-39-05) using Lipofectamine 2000 (ThermoFisher Scientific, Cat #11668027). Medium was replaced four hours after stimulation with DMEM 10% FCS. Activities of the luciferases were measured 24 h after stimulation.

To investigate the IFN-I response pathway, ninety-six-well plates were coated with 50 μg/mL of poly-D-lysine (ThermoFisher Scientific, Cat# A3890401) for 30 min at 37°C, then washed with water and dried before being seeded with 4 × 10^4^ HEK293T cells per well. Each well was transfected the day after with 12.5 ng of pISRE-Luc, 5 ng of phRL-TK, together with 35.4 ng of plasmids expressing the viral proteins or the mCherry control, using JetOptimus reagent (Ozyme, Cat# POL101000006) according to the manufacturer’s recommendations. Cells were treated the day after with 1000 U/mL of IFNα-2a (ImmunoTools, Cat#11343506) for 18 h before measuring the activity of the luciferases.

To monitor the NF-κB pathway, 2×10^4^ HeLa cells per well were seeded in ninety-six-well plates. Each well was transfected the day after with 12.5 ng of pSGNluc, 6.25 ng of phRL-TK, together with 31.25 ng of plasmids expressing the viral proteins or the mCherry control, using JetOptimus reagent (Ozyme, Cat# POL101000006) according to the manufacturer’s recommendations. Cells were treated 24 h later with 10 ng/μL of human IL-1β (Peprotech, Cat# 200-01B) for 2 h before measuring the luciferase activities.

For all assays, the activities of the luciferases were measured using the Dual-Glo Luciferase Assay system according to the manufacturer’s protocol (Promega, Cat# E2920) and a microplate reader (Tristar 5, Berthold) with acquisition for 1 s. For each well, the relative luciferase activity was calculated as the ratio between the Firefly and Renilla luciferase activities. For each condition, experiments were performed in triplicate and repeated in four separate experiments.

### Immunofluorescence

For the detection of viral proteins, 2×10^5^ HeLa cells were seeded on 12 mm glass coverslips placed in a 12-well plate. Twenty-four hours later, cells were transfected with 0.5 μg of plasmid and fixed 48 h after transfection with 4% paraformaldehyde. Cells were permeabilised with 0.1% Triton X-100 in PBS for 20 min and blocked for 30 min in PBS - 2% BSA. Staining was done overnight at 4°C using mouse anti-Flag antibody (Merck, Cat# F1804) diluted in PBS – 0.5% BSA, followed by incubation for 1 h at room temperature with DAPI (Thermo FisherTM, Cat# 62248) and Alexa Fluor 488 goat anti-mouse antibody (ThermoFisher Scientific) diluted at 1.3 μg/mL in the same buffer.

For the detection of p65, 2×10^5^ HeLa cells were seeded on 12 mm glass coverslips placed in a 12-well plate and transfected 24 h later with 2 μg of plasmid expressing NiV or LayV W or W_CTD_ proteins. Twenty-four hours after transfection, cells were stimulated or not with 10 ng/mL of human IL-1β (Peprotech, Cat# 200-01B) for 20 min at 37°C before being washed with ice-cold PBS and fixed in 4% paraformaldehyde for 30 min on ice. Cells were then permeabilised for 10 min in TBS – 0.1% Triton X-100 and blocked for 10 additional minutes in TBS – 2.5% BSA. Then, cells were stained overnight at 4°C with rabbit anti-NF-κB p65 (Cell Signalling, Cat#8242) and mouse anti-Flag antibody (Merck, Cat# F1804) diluted in TBS– 2% FCS - 0.1% Tween. The day after, cells were incubated for 1 h at room temperature with DAPI (ThermoFisher Scientific, Cat# 62248) together with Alexa Fluor 488 donkey anti-rabbit (ThermoFisher Scientific, Cat# A21206) and Alexa Fluor 555 goat anti-mouse antibodies (ThermoFisher Scientific, Cat# A21422) diluted at 1.3 μg/mL in the same buffer.

Coverslips were mounted with Fluoromount-G (SouthernBiotech, Cat# 0100-01) before observation on a confocal microscope (Zeiss, LSM800Airy Scan) hosted in the PLATIM microscopy facility (SFR Biosciences, Lyon, France). Images were treated using ImageJ software.

### Quantification and statistical analysis

#### Luciferase assay analysis

All assays with statistical analysis were performed in four independent experiments, with each condition tested in triplicate (three technical replicates per plate). For each well, the Firefly luciferase signal was normalised by the Renilla luciferase signal. Statistical analyses were done using the non-parametric Mann-Whitney test.

#### Image quantification

Fluorescence microscopy images were analysed and quantified using QuPath 0.5.1 software (https://qupath.github.io/). Cells were identified using a cell detection program based on the staining of the nuclei (DAPI staining). For the quantification of the NF-κB p65 signal, the nuclear signal is defined as the signal present in the mask created by the DAPI staining, and the cytoplasmic signal as the signal present in a 2 µm radius outside of the nucleus mask. Each condition was tested in four independent experiments with three images analysed per experiment, leading to a total of 150 cells analysed per condition. Statistical analysis was done using the non-parametric Mann-Whitney test.

#### Phylogenetic tree

The L protein sequences from Nipah virus (NC_002728), Hendra virus (NC_001906), Cedar virus (NC_025351), Ghana virus (NC_025256), Yunnan bat henipavirus 1 (PQ621839), Yunnan bat henipavirus 2 (PQ621840), Angavokely henipavirus (ON613535), Gamak virus (MZ574407), Crocidura tanakae henipavirus 2 (PP272531), Jingmen Crocidura shantungensis henipavirus 1 (OM030314), Wufeng Crocidura attenuata henipavirus 1 (OM030317), Crocidura tanakae henipavirus 1 (OQ970176), Resua virus (OR713876), Wufeng Chodsigoa smithii henipavirus 1 (OM030316), Chodsigoa hypsibia henipavirus (OQ236120), Camp Hill virus (PQ140948), Ninorex virus (OQ438286), Denwin virus (OR713883), Crocidura olivieri henipavirus 2 (PQ541139), Melian virus (OK623353), Jingmen Crocidura shantungensis henipavirus 2 (PP272750), Daeryong virus (MZ574409), Crocidura olivieri henipavirus 1 (PQ541138), Lechcodon virus (OR713879), Wenzhou shrew henipavirus 1 (OQ715593), Hasua virus (OR713881), Mojiang virus (NC_025352), and Langya virus (OM101125), were used to build a multiple sequence alignment using MAFFT [42]. The phylogenetic tree was calculated with the Neighbor Joining algorithm in MAFFT.

## RESULTS

### Similar intracellular localisation of NiV and LayV P and V proteins, but different localisation of their W proteins

While NiV P and V proteins are found in the cytoplasm, W is concentrated in the nucleus [29,43,44,37]. To investigate if LayV proteins harbour the same cellular localisation, we transfected NiV and LayV proteins fused to a Flag tag in HeLa cells and observed their localisations by immunofluorescence. Similar to NiV, LayV P and V proteins are both localised in the cytoplasm (Figure 2A, B). On the contrary, NiV and LayV W proteins are found in different cell compartments, with NiV W being restricted to the nucleus while LayV W is diffused in the cytoplasm (Figure 2C). These results correlate with the presence of a nuclear localisation signal (NLS) located on NiV W_CTD_ and the absence of conservation of these residues in LayV W (Figure S1D) [45].

**Figure 2.**
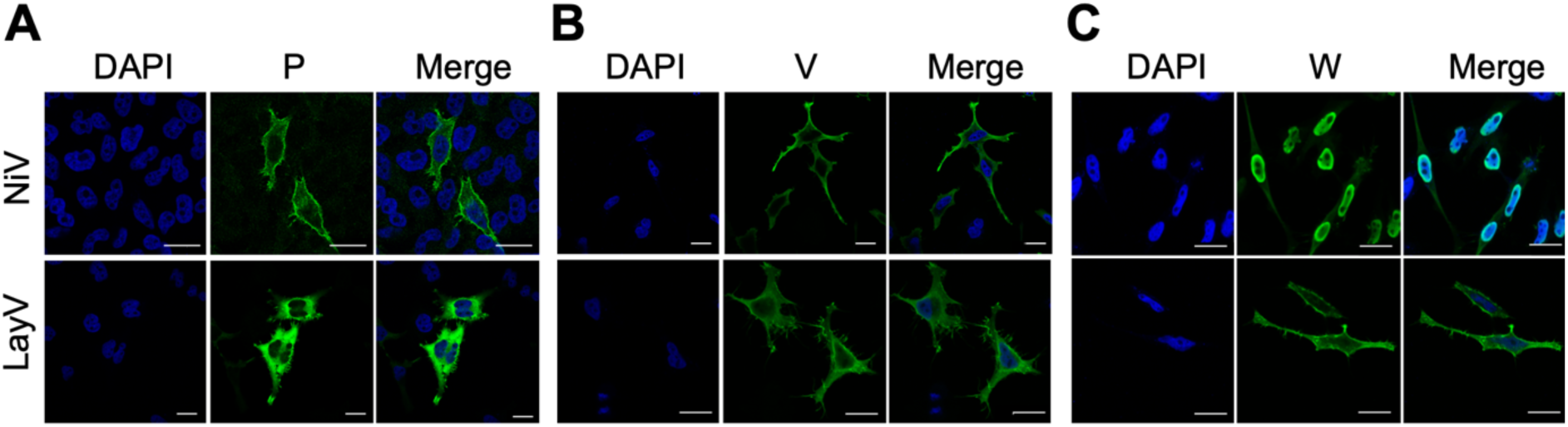
Localisation of NiV and LayV P, V, and W proteins. HeLa cells were transfected to express NiV or LayV Flag-tagged P (A), V (B), or W (C). At 24 h.p.t., proteins were stained using an anti-Flag antibody and imaged by confocal microscopy. Viral proteins are in green, and nuclei are in blue. Scale bars correspond to 20 µm.

### NiV and LayV P proteins inhibit the IFN-I signalling pathway

Via their common N-terminal domain (P_NTD_), NiV P, V, and W proteins inhibit the transcription factor STAT1, leading to a decrease in IFN-I signalling [27–31,26,32,33]. To monitor this effect, the highly transfectable HEK293T cells were transfected with plasmids encoding the viral proteins and a reporter gene under the control of the Interferon-Stimulated Response Element (ISRE). After stimulation with IFN-2α, the reporter signal was significantly decreased for cells expressing NiV and LayV P proteins compared to the control cells expressing the fluorescent protein mCherry (Figure 3A). Similarly to NiV, LayV P_NTD_ is sufficient to mediate this inhibition (Figure 3B). However, although LayV P and P_NTD_ expression levels were higher or similar to NiV proteins, LayV proteins were less efficient in blocking the signalling (Figure 3, S2A). This difference may be linked to the low conservation of the STAT1-binding region identified on NiV P and thus to a lower affinity for human STAT1 (Figure S1A).

**Figure 3.**
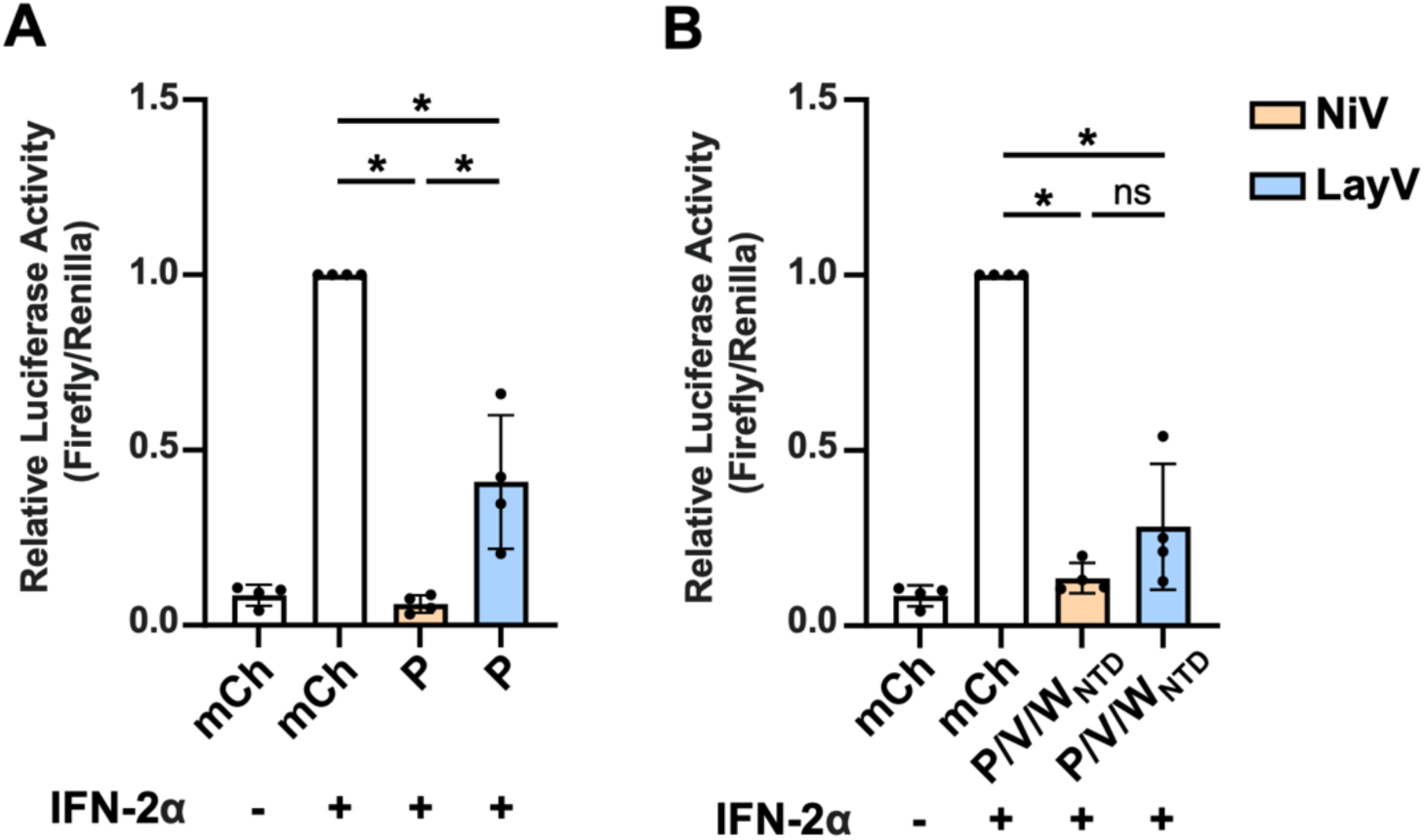
NiV and LayV P/V/W_NTD_ inhibit the IFN-I signalling pathway. HEK293T cells were co-transfected with three plasmids coding for (1) a viral protein or the mCherry (mCh) control, (2) the Firefly luciferase under an ISRE-dependent promoter, and (3) the Renilla luciferase under a constitutive promoter. Either the full-length P proteins are expressed (A) or only the N-terminal domain common to P, V, and W (P/V/W_NTD_) (B). At 18 h.p.t., cells were stimulated with 1000 U/mL IFN-2α for 24 h before measuring the luciferase activities. Mean and SD were calculated from four independent experiments.

### NiV and LayV V proteins inhibit MDA5 but not RIG-I signalling pathways

The C-terminal domain of V proteins is capable of binding MDA5, disrupting its folding, and inhibiting the subsequent activation of the IFN-β promoter [34–36,46]. To compare the effect of NiV and LayV V proteins on the IFN-β induction pathway after MDA5 or RIG-I transfection and activation, we used Huh-7.5 cells since they do not express RIG-I and only a low level of MDA5 [38]. NiV and LayV V proteins are both capable of inhibiting the induction of the IFN-β promoter upon MDA5 activation (Figure 4A, S2B). This effect is even stronger when only the V_CTD_ is expressed as a fusion protein with mCherry (mCh-V_CTD_) (Figure 4B, S2B). As expected, a mutant of NiV V_CTD_ carrying the mutations E411A and I414A, known to impair the binding to MDA5 [47], is no longer able to inhibit the activation of the IFN-β promoter (Figure 4B). On the contrary, neither NiV nor LayV V proteins nor the mCh-V_CTD_ constructs inhibited the activation of the IFN-β promoter after RIG-I stimulation (Figure 4C, D).

**Figure 4.**
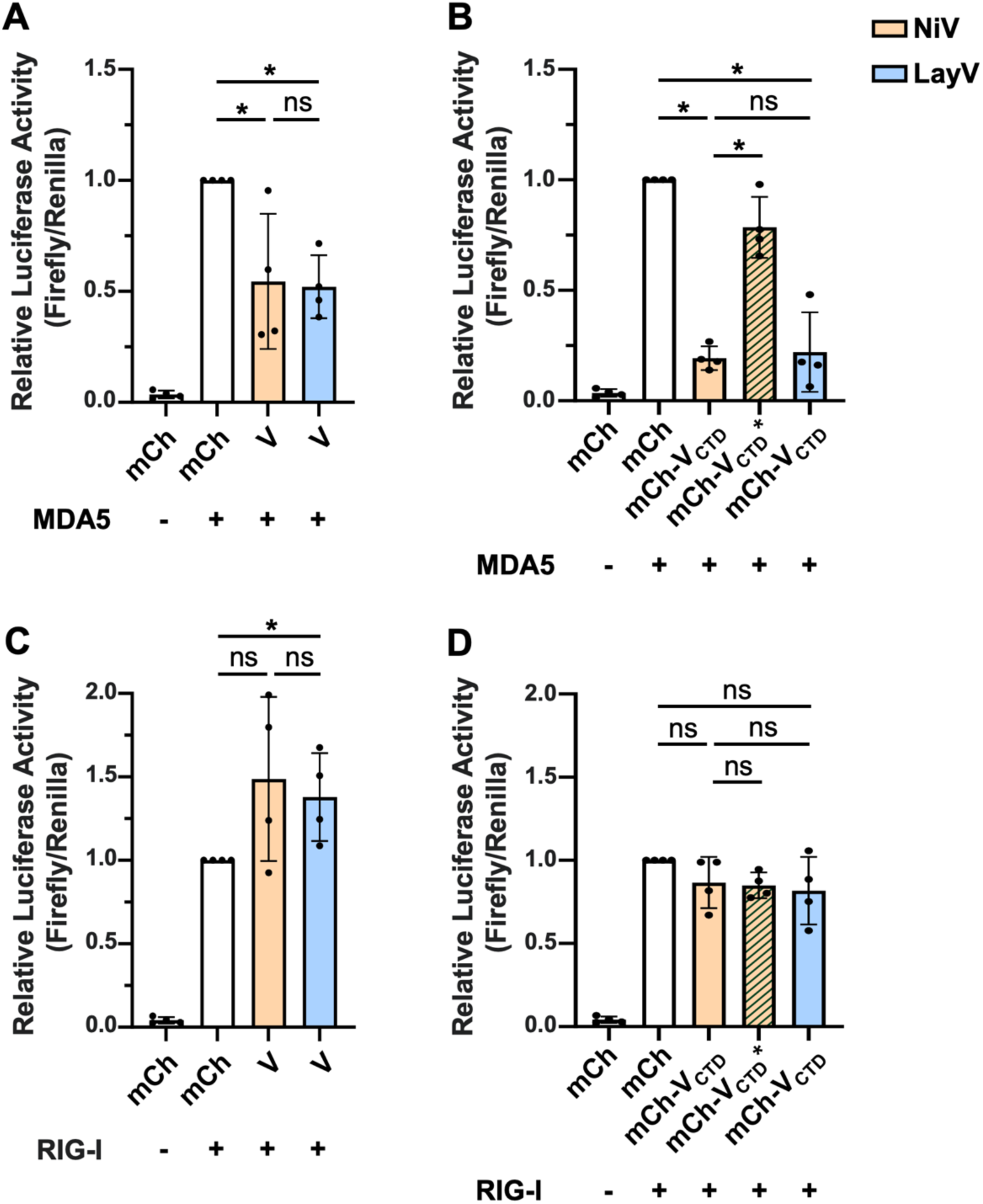
NiV and LayV V proteins inhibit MDA5 but not RIG-I signalling pathway. Huh-7.5 cells were co-transfected with four plasmids coding for (1) human MDA5 (A, B) or RIG-I (C, D), (2) a viral protein or the mCherry control, (3) the Firefly luciferase under the IFN-β promoter, and (4) the Renilla luciferase under a constitutive promoter. Viral proteins are expressed either as full-length proteins (A, C) or as fusion proteins between mCherry and the V C-terminal domain (mCh-V_CTD_) (B, D). Cells were stimulated 24 h.p.t. by transfecting 100 ng Poly(I:C), and the luciferase activities were measured 48 h.p.t. Mean and SD were calculated from four independent experiments. mCh-V_CTD_* carries the mutations E411A and I414A.

Overall, these results indicate that both NiV and LayV V can inhibit the IFN-β induction pathway after the activation of MDA5 but not RIG-I.

### The W protein of NiV, but not LayV, inhibits MDA5 and RIG-I signalling pathways

When V proteins were replaced with W in the MDA5 and RIG-I activation assays, NiV W was able to inhibit the activation of the IFN-β promoter upon both MDA5 (Figure 5A) and RIG-I (Figure 5C) activation. This effect was stronger when only the CTD of NiV W protein was expressed (mCh-W_CTD_) (Figure 5B, D). Conversely, LayV W protein did not have any inhibitory effect on the activation of the IFN-β promoter upon MDA5 and RIG-I activation (Figure 5). To compensate for the lower expression of LayV W compared to NiV W (Figure S2C), we transfected a range of plasmid quantities. While we detected an inverse correlation between the amount of NiV W and the reporter signal intensity, the activation of the reporter’s promoter was insensitive to the increasing amount of LayV W, even when expressed at higher levels than NiV W, thus indicating that the difference in inhibitory effect is protein-specific and not due to the protein expression levels (Figure S3). This is further confirmed with the mCh-W_CTD_ constructs, which are similarly expressed, with only NiV W_CTD_ being able to inhibit the induction (Figure 5B, D, S2C).

**Figure 5.**
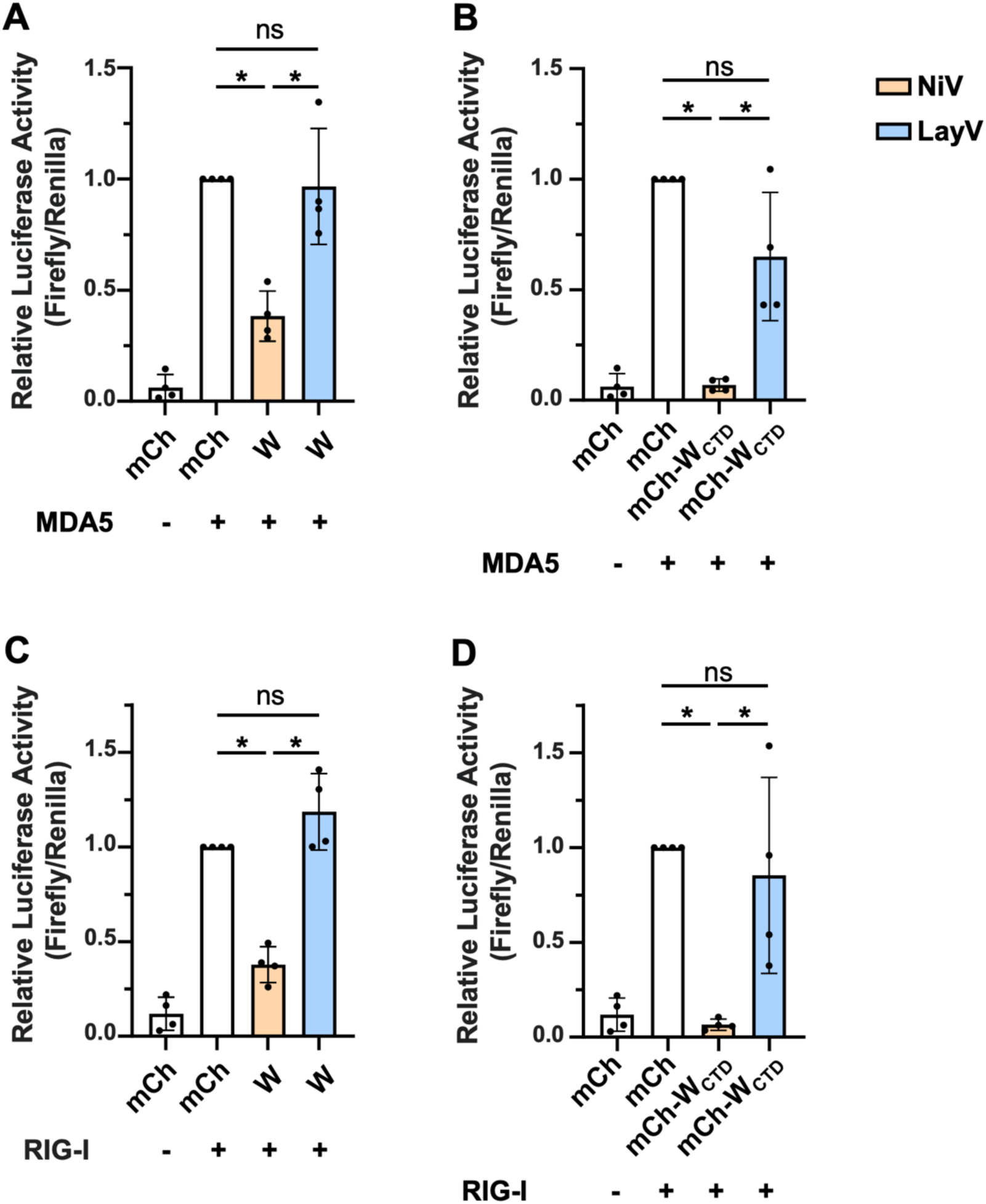
Only NiV W protein inhibits MDA5 and RIG-I signalling pathways. Huh-7.5 cells were co-transfected with four plasmids coding for (1) human MDA5 (A, B) or RIG-I (C, D), (2) a viral protein, (3) the Firefly luciferase under the IFN-β promoter, and (4) the Renilla luciferase under a constitutive promoter. Viral proteins are expressed either as full-length proteins (A, C) or as fusion proteins between mCherry and the W C-terminal domain (mCh-W_CTD_) (B, D). Cells were stimulated 24 h.p.t. by transfecting 100 ng of Poly(I:C), and luciferase activities were measured 48 h.p.t. Mean and SD were calculated from four independent experiments.

While LayV W is found in the cytoplasm, NiV W is concentrated in the nucleus thanks to an NLS present on W_CTD_ (Figure 1B, S1D). Since this signal is required for NiV W inhibitory activity of immune pathways [37,43], we examined whether the absence of inhibitory activity of LayV W would be due to its cytoplasmic localisation. Mutating NiV W NLS (NLSko) relocated the mCh-W_CTD_ fusion protein in the cytoplasm, and adding back the exogenous NLS of the simian vacuolating virus 40 (SV40_NLS_) partially restored the nuclear localisation (Figure 6A). As expected, the inhibitory effect of these NiV mCh-W_CTD_ constructs on the IFN-β promoter correlated with their intracellular localisation (Figure 6A, B). On the contrary, while the addition of the SV40_NLS_ to LayV mCh-W_CTD_ also partially relocated the protein to the nucleus, this relocalisation did not provide any significant inhibitory effect to LayV W_CTD_ (Figure 6A, C). Thus, the absence of an inhibitory effect of LayV W is not solely due to its cytoplasmic localisation.

**Figure 6.**
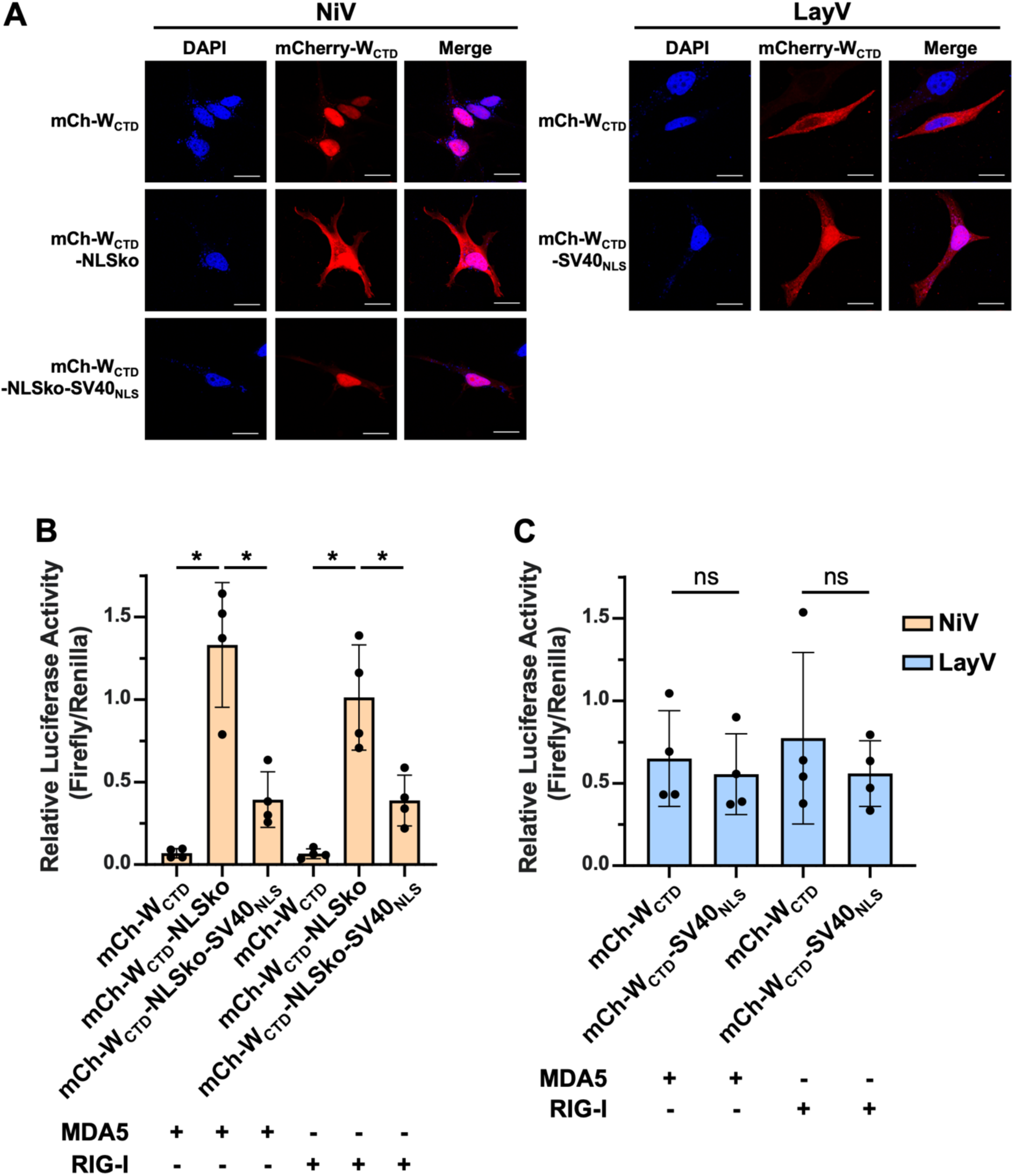
The cytoplasmic localisation of LayV W is not responsible for the absence of inhibition of the MDA5 and RIG-I pathways. (A) HeLa cells were transfected to express fusion proteins between mCherry and NiV or LayV W C-terminal domain (mCh-W_CTD_). In some constructs, the nuclear localisation signal (NLS) of NiV W was mutated (NLSko), and the NLS from SV40 was added to NiV and LayV W_CTD_ (SV40_NLS_). Twenty-four hours post-transfection, stained cells were imaged by confocal microscopy. Viral proteins are in red and nuclei in blue. Scale bars correspond to 20 µm. (B, C) Huh-7.5 cells were co-transfected with four plasmids coding for (1) human MDA5 or RIG-I, (2) a viral protein or the mCherry control, (3) the Firefly luciferase under the IFN-β promoter, and (4) the Renilla luciferase under a constitutive promoter. Cells were stimulated 24 h.p.t. by transfecting 100 ng of Poly(I:C), and relative activities of the luciferases were measured 48 h.p.t. Mean and SD were calculated from four independent experiments.

### The NF-κB pathway and the translocation of p65 into the nucleus are impaired by NiV but not LayV W protein

We have previously demonstrated that NiV W can inhibit the NF-κB pathway via its CTD [37]. Here, we analysed the effect of LayV and NiV proteins on the NF-κB pathway after stimulation of cells with IL-1β (Figure 7A, B). Although weakly for the full-length proteins, the expression of NiV W and W_CTD_ decreased the activation of the NF-κB promoter, while LayV proteins did not and even increased the activity (Figure 7A, B).

**Figure 7.**
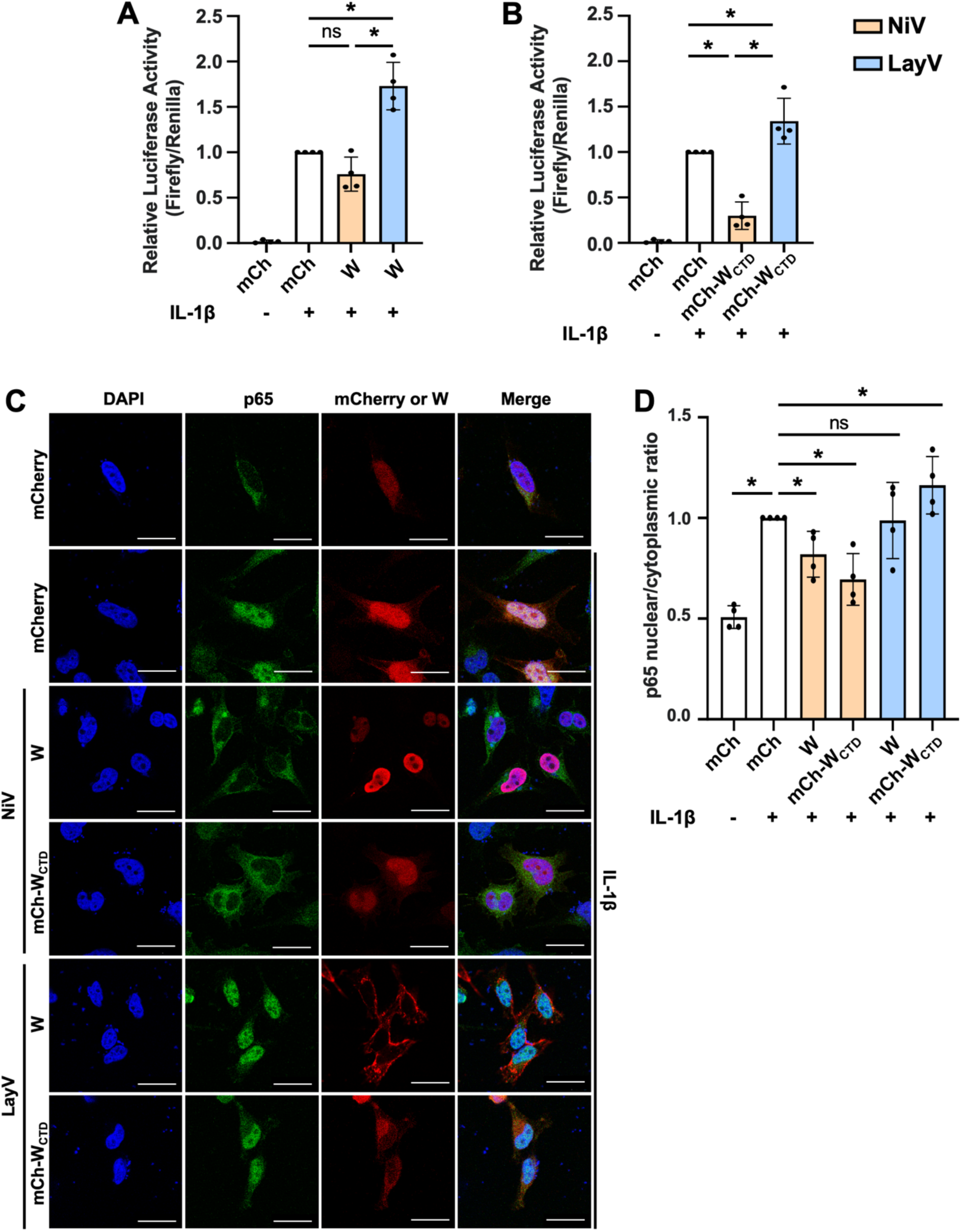
NiV but not LayV W protein can block the NF-κB pathway. (A and B) HeLa cells were co-transfected with three plasmids coding for (1) a viral protein or the mCherry control, (2) the Firefly luciferase under an NF-κB-dependent promoter, and (3) the Renilla luciferase under a constitutive promoter. Viral proteins are expressed either as full-length proteins (A) or as fusion proteins between mCherry and the W C-terminal domain (mCh-W_CTD_) (B). At 18 h.p.t., cells were incubated with 10 ng/mL IL-1β for 2 h before measuring the luciferase activities. Mean and SD were calculated from four independent experiments. (C) HeLa cells were transfected to express NiV or LayV W or mCh-W_CTD_ viral proteins. At 24 h.p.t., cells were stimulated with 10 ng/mL IL-1β for 20 min before being fixed. W was stained with an anti-Flag antibody and endogenous p65 protein with an anti-NF-κB p65 antibody. Stained cells were imaged by confocal microscopy. Nuclei are in blue, p65 in green, and viral proteins in red. Scale bars correspond to 20 µm. (D) The mean intensity of nuclear and cytoplasmic NF-κB p65 was evaluated using QuPath software on an average of 150 events per condition. Mean and SD were calculated from four independent experiments.

Since NiV W decreases the nuclear concentration of NF-κB p65 subunit in infected cells, we monitor the localisation of p65 after NiV or LayV W expression and IL-1β stimulation (Figure 7C, D). While NiV W decreased the concentration of p65 in the nucleus via its CTD, LayV W did not prevent the relocalisation of p65 in the nucleus upon IL-1β stimulation. These results suggest that, unlike NiV W, LayV W is not able to inhibit the accumulation of p65 in the nucleus and thus the subsequent activation of the NF-κB promoter.

## DISCUSSION

The presence of the entry receptor(s) and the ability to counteract the host defence are among the most important barriers for a virus to successfully infect a new host. For paramyxoviruses, the proteins translated from the P gene (P, V, W, and C) are the main anti-host defence factors. In the absence of a reverse genetic system for Langya virus, we used cell transfection assays to assess the ability of LayV P, V, and W proteins to block several pathways of the intrinsic immunity in human cells and compared them to NiV proteins, a closely related, zoonotic, and highly pathogenic virus.

We report that both LayV and NiV V proteins can efficiently inhibit transcription activation after MDA5 stimulation. Since the amino acids essential for the folding of the V_CTD_ and the binding to MDA5 are highly conserved between NiV and LayV, and among henipa- and parahenipaviruses (Figure S1C), LayV V likely directly binds to and blocks the activation of MDA5. Although NiV V was described to decrease RIG-I activation via the inhibition of the TRIM25-mediated ubiquitination of RIG-I [48], we did not observe any effect in our assays. This discrepancy likely reflects differences in the assays, mainly the absence of overexpression of TRIM25 and the use of different cell lines (HEK293T vs Huh-7.5). Thus, we cannot conclude on the effect of LayV V on the TRIM25-mediated ubiquitination of RIG-I.

Similarly, LayV P can inhibit the signalling under the IFN-I receptor, but less efficiently than NiV P. This difference does not seem correlated with P expression levels and may reflect differences in their ability to bind human STAT proteins, as suggested by the rather low conservation of the STAT1-binding region of NiV P.

Unlike NiV W, LayV W was unable to inhibit transcription activation after activation of MDA5, RIG-I, and the IL-1β receptor. Since the transcription factor NF-κB is activated by each of these signalling pathways, and is known to be inhibited by NiV W, NiV W may inhibit these three pathways using the same molecular mechanism: recruiting the cellular protein 14-3-3 to the nucleus to favour the export of p65 [37,49]. The relocation of LayV W to the nucleus, where NiV W is concentrated, did not provide any functional effect, and LayV W lacks the residues of NiV W known to bind 14-3-3. Thus, unlike NiV W, LayV W is most likely not able to recruit 14-3-3 to the nucleus to inhibit p65 accumulation.

Since the C protein also participates in the immune evasion, and LayV is thought to express an additional protein upstream of the F open reading frame (Figure 1B), these two proteins may contribute to the immune evasion and compensate for the absence of activity of LayV W in inhibiting the human immunity pathways.

Overall, this study suggests that the P and W proteins of LayV are not as efficient as NiV proteins in counteracting human intrinsic immunity. Thus, since the inhibition of the innate immune system is a major factor of pathogenicity, this distinction between NiV and LayV may be one of the reasons for the difference in pathogenicity observed in humans, with the high lethality of NiV-infected patients, in contrast to LayV infection.

## ACKNOWLEDGEMENTS

We acknowledge the contribution of the SFR Biosciences (Université Claude Bernard Lyon 1, CNRS UAR3444, Inserm US8, ENS de Lyon) LYMIC-PLATIM microscopy platform for training and the use of the LSM800 AS confocal microscope. We thank Mathieu Iampietro and Lucia Amurri (CIRI, Lyon) for their advice on the NF-κB experiments and all current and former members of the team Immunobiology of Viral Infections at CIRI for their help in the realisation of the study. We thank Omran Allatif (CIRI, bioinformatics and biostatistics (BIBS) platform) for his help with statistical analysis.

## DATA AVAILABILITY STATEMENT

Data is available upon request.

