## Supplementary Material for "Differential Immunomodulatory Properties of Langya and Nipah Virus Proteins"

#### MATERIALS AND METHODS

##### *Protein alignments*

Sequences from Nipah virus (NC_002728), Hendra virus (NC_001906), Cedar virus (NC_025351), Ghana virus (NC_025256), Yunnan bat henipavirus 1 (PQ621839), Yunnan bat henipavirus 2 (PQ621840), Angavokely henipavirus (ON613535), Gamak virus (MZ574407), Crocidura tanakae henipavirus 2 (PP272531), Jingmen Crocidura shantungensis henipavirus 1 (OM030314), Wufeng Crocidura attenuata henipavirus 1 (OM030317), Crocidura tanakae henipavirus 1 (OQ970176), Resua virus (OR713876), Wufeng Chodsigoa smithii henipavirus 1 (OM030316), Chodsigoa hypsibia henipavirus (OQ236120), Camp Hill virus (PQ140948), Ninorex virus (OQ438286), Denwin virus (OR713883), Crocidura olivieri henipavirus 2 (PQ541139), Melian virus (OK623353), Jingmen Crocidura shantungensis henipavirus 2 (PP272750), Daeryong virus (MZ574409), Crocidura olivieri henipavirus 1 (PQ541138), Lechcodon virus (OR713879), Wenzhou shrew henipavirus 1 (OQ715593), Hasua virus (OR713881), Mojiang virus (NC_025352), and Langya virus (OM101125), were used to build multiple sequence alignment of P, $V_{CTD}$, and $W_{CTD}$ using MAFFT (gap opening penalty = 5) [1]. The alignments are displayed using Jalview and the Clustal colour code [2].

##### *Western blot*

Transfected HeLa cells were lysed 24 h after transfection in ice-cold RIPA lysis buffer (Life Technologies, Cat# 89901) supplemented with Halt protease inhibitor cocktail (ThermoFisher Scientific, Cat# 78429) for 30 min on ice and then centrifuged at 11,000 g for 15 min. The protein concentrations of the lysates were quantified using Pierce BCA Protein assay

(ThermoFisher Scientific, Cat# 23227). Lysates were denatured in SDS Laemmli buffer containing 1% β-mercaptoethanol and heated for 10 min at 96°C. Identical total protein quantities of each sample were then separated on TGX protein gels (Biorad, Cat# 4569035) and transferred onto polyvinylidenedifluoride (PVDF) membranes (Biorad, Cat# 1704157). PVDF membranes were blocked in phosphate-buffered saline solution containing 5% milk for 30 min at room temperature and then incubated overnight at 4°C with primary antibodies, mouse anti-Flag (Merck, Cat# F1804), mouse anti-GAPDH (Chemicon, Cat# MAB374), or rabbit anti-vinculin (Invitrogen, Cat# 700062) diluted in PBS - 0.1% Tween20 - 0.5% milk. After washing with PBS - 0.1% Tween, membranes were incubated for 1 h at room temperature with horseradish peroxidase-conjugated mouse (Promega, Cat# W4021) or rabbit (Promega, Cat# W4011) secondary antibodies diluted in PBS - 0.1% Tween20 - 0.5% milk. Secondary antibodies were revealed with the Super Signal West Dura reagent (ThermoScientific™, Cat# 34076), and chemiluminescent signals were measured with ImageQuant LAS500 (GE Healthcare).

### SUPPLEMENTARY FIGURES

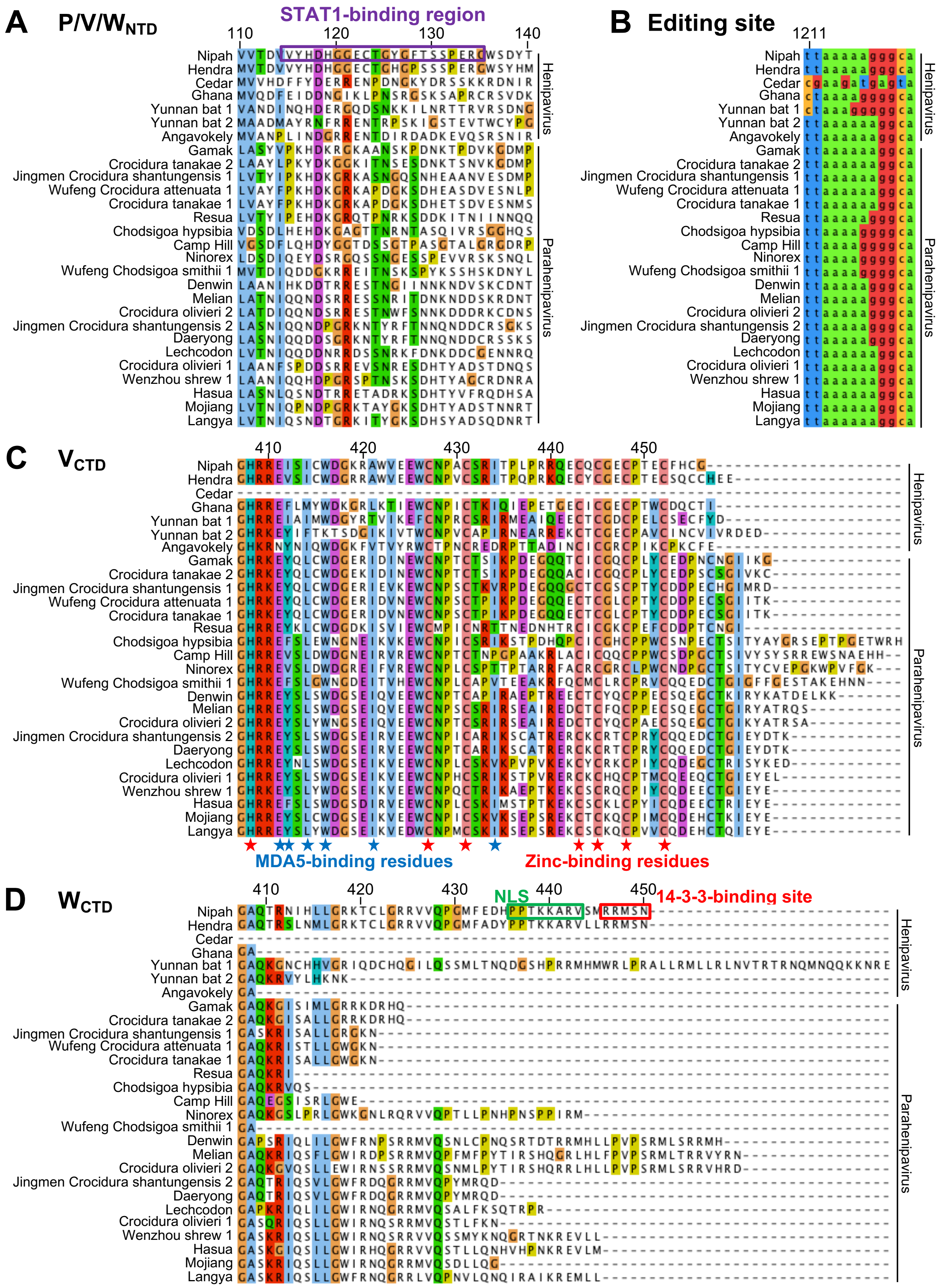

***Supplementary Figure 1. Alignment of henipavirus and parahenipavirus sequences.***

Alignment of the protein sequences of a fragment of NiV P/V/$W_{NTD}$ (A), the nucleotide sequence of the editing site (B), and the protein sequences of the $V_{CTD}$ (C) and the $W_{CTD}$ (D) of henipaviruses and parahenipaviruses. The STAT1-binding region identified on NiV is shown with a purple rectangle. MDA5-binding residues and zinc-binding residues are shown with blue and red stars, respectively. NiV W NLS and 14-3-3-binding site are shown in a green and red rectangle, respectively.

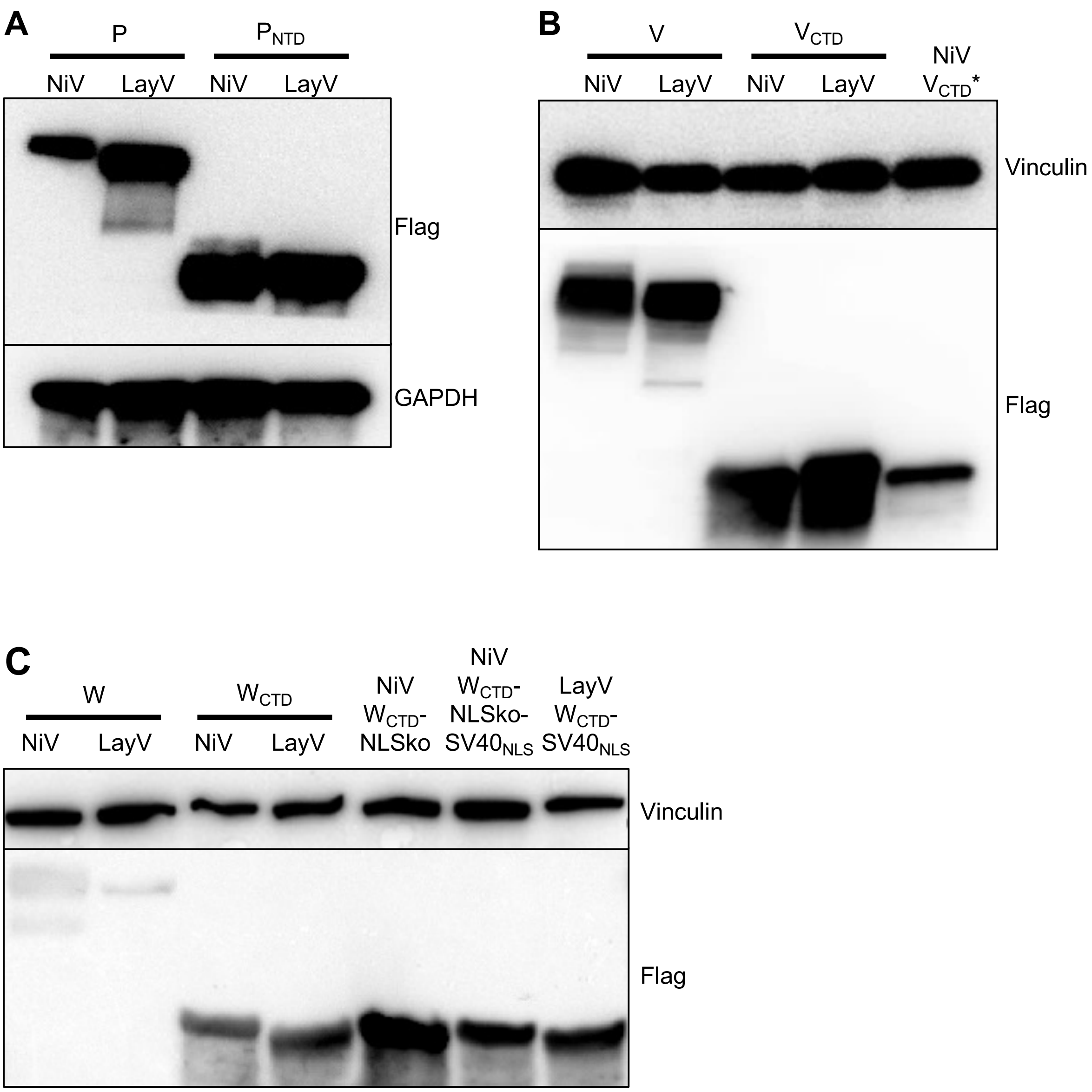


***Supplementary Figure 2. Expression of NiV and LayV P, V, and W proteins.***

HeLa cells were transfected with plasmids coding for NiV and LayV full-length P or P N-terminal domain ($P_{NTD}$) (A), full-length V or V C-terminal domain ($V_{CTD}$) (B), or full-length W or W C-terminal domain ($W_{CTD}$) (C). At 48 h.p.t., cells were harvested, and the protein expression was analysed by SDS-PAGE and western blot using anti-Flag antibody. GAPDH and vinculin are used as housekeeping controls.

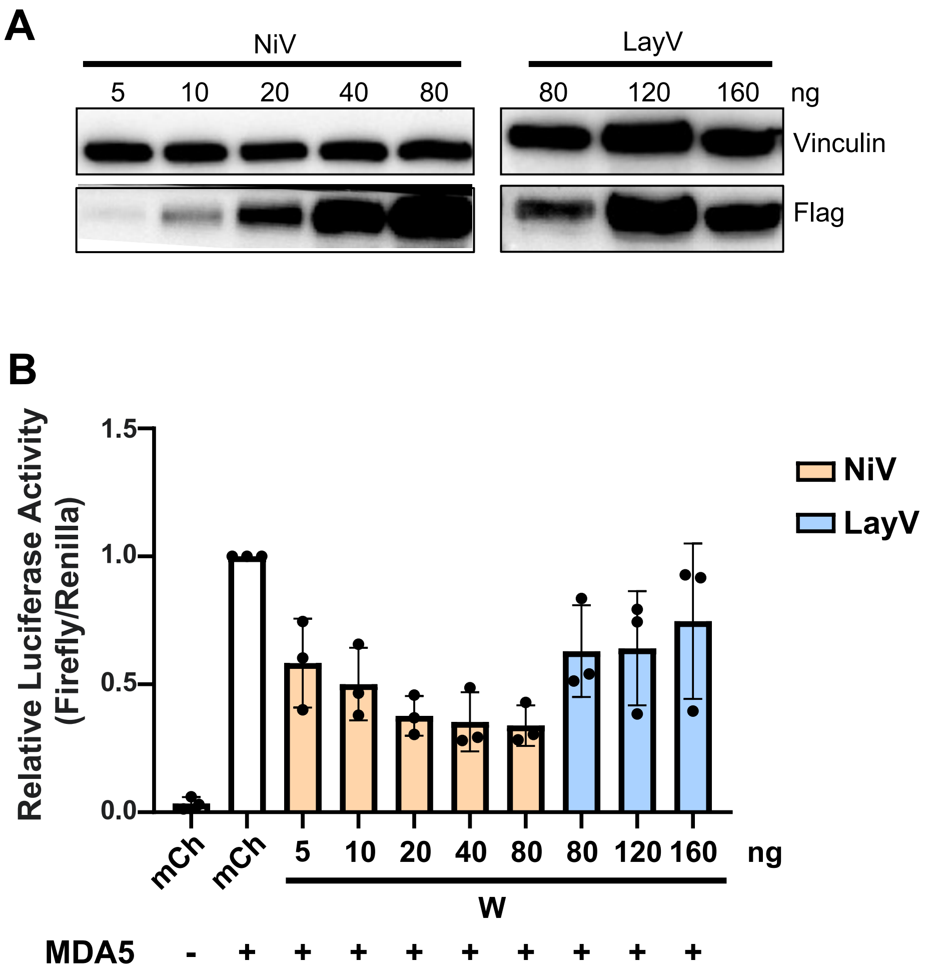


***Supplementary Figure 3. The absence of an inhibitory effect of LayV W on the MDA5 pathway is not due to lower protein expression.***

(A) HeLa cells were transfected with increasing amounts of plasmids coding for NiV and LayV full-length W proteins. At 48 h.p.t., cells were lysed, and protein expression was analysed by SDS-PAGE and western blot using anti-Flag antibody. Vinculin is used as a housekeeping control. (B) Huh-7.5 cells were co-transfected with four plasmids coding for (1) human MDA5, (2) NiV or LayV W protein, (3) the Firefly luciferase under the interferon beta promoter, and (4) the Renilla luciferase under a constitutive promoter. Cells were stimulated the day after by transfecting 100 ng Poly(I:C), and luciferase activities were measured 48 h.p.t. Mean and SD were calculated from three independent experiments.
